# Tokenized and Continuous Embedding Compressions of Protein Sequence and Structure

**DOI:** 10.1101/2024.08.06.606920

**Authors:** Amy X. Lu, Wilson Yan, Kevin K. Yang, Vladimir Gligorijevic, Kyunghyun Cho, Pieter Abbeel, Richard Bonneau, Nathan Frey

## Abstract

Existing protein machine learning representations typically model either the sequence or structure distribution, with the other modality implicit. The latent space of sequence-to-structure prediction models such as ESMFold represents the *joint distribution* of sequence and structure; however, we find these embeddings to exhibit massive activations, whereby some channels have values 3000× higher than others, regardless of the input. Further, on continuous compression schemes, ESMFold embeddings can be reduced by a factor of 128× along the channel and 8× along the length, while retaining structure information at <2Å scale accuracy, and performing competitively on protein function and localization benchmarks. On discrete compression schemes, we construct a tokenized all-atom structure vocabulary that retains high reconstruction accuracy, thus introducing a *tokenized representation of all-atom structure that can be obtained from sequence alone*. We term this series of embeddings as CHEAP (Compressed Hourglass Embedding Adaptations of Proteins) embeddings, obtained via the HPCT (Hourglass Protein Compression Transformer) architecture. CHEAP is a compact representation of both protein structure and sequence, sheds light on information content asymmetries between sequence and structure, democratizes representations captured by large models, and is designed to have flexible downstream applications such as generation, search, and prediction.

## 1. Introduction

Structure is an important determinant in scaffolding functions and biomolecular interactions. Real-world manufacturing of proteins require precise specification of sequence, and most extant functional biomolecules result from evolution in sequence space. Capturing useful information from all available data for proteins may unlock powerful new capabilities for protein design. Protein sequence datasets can be 10^2^ to 10^4^ times larger than structural datasets. As of 2024, structural datasets such as the PDB [2] contains 218,196 samples, while sequence-only datasets such as UniRef50 [38] contains 63,849,054 samples. Being able to accurately capture high-precision structural information from sequence input is therefore desirable for machine learning applications in biology.

In recent years, sequence-to-structure prediction [20, 24] or structure-to-sequence design [6, 18] models have become increasingly capable. Such models have been described as “protein structure foundation models” [44], mirroring a broader paradigm in AI where large models pretrained on large corpora of data can be adapted for an array of downstream tasks [4]. These model capture intricate information that emerges with scale, but in their native formulation as prediction models, they are unable to perform generation, similarity search, property prediction, etc., thus limiting their use as a foundation model.

Since sequence-to-structure models map from *p*(sequence) to *p*(structure), its intermediate latent space can be viewed as a representation of the joint distribution *p*(sequence, structure) (Figure 1B). Of particular interest, ESMFold [24] demonstrates that sequence-to-structure prediction can be built on top of protein language model (pLM) embeddings (Figure 1A). This provides a compact latent space of joint structure and sequence information (Figure 1C).

**Figure 1:**
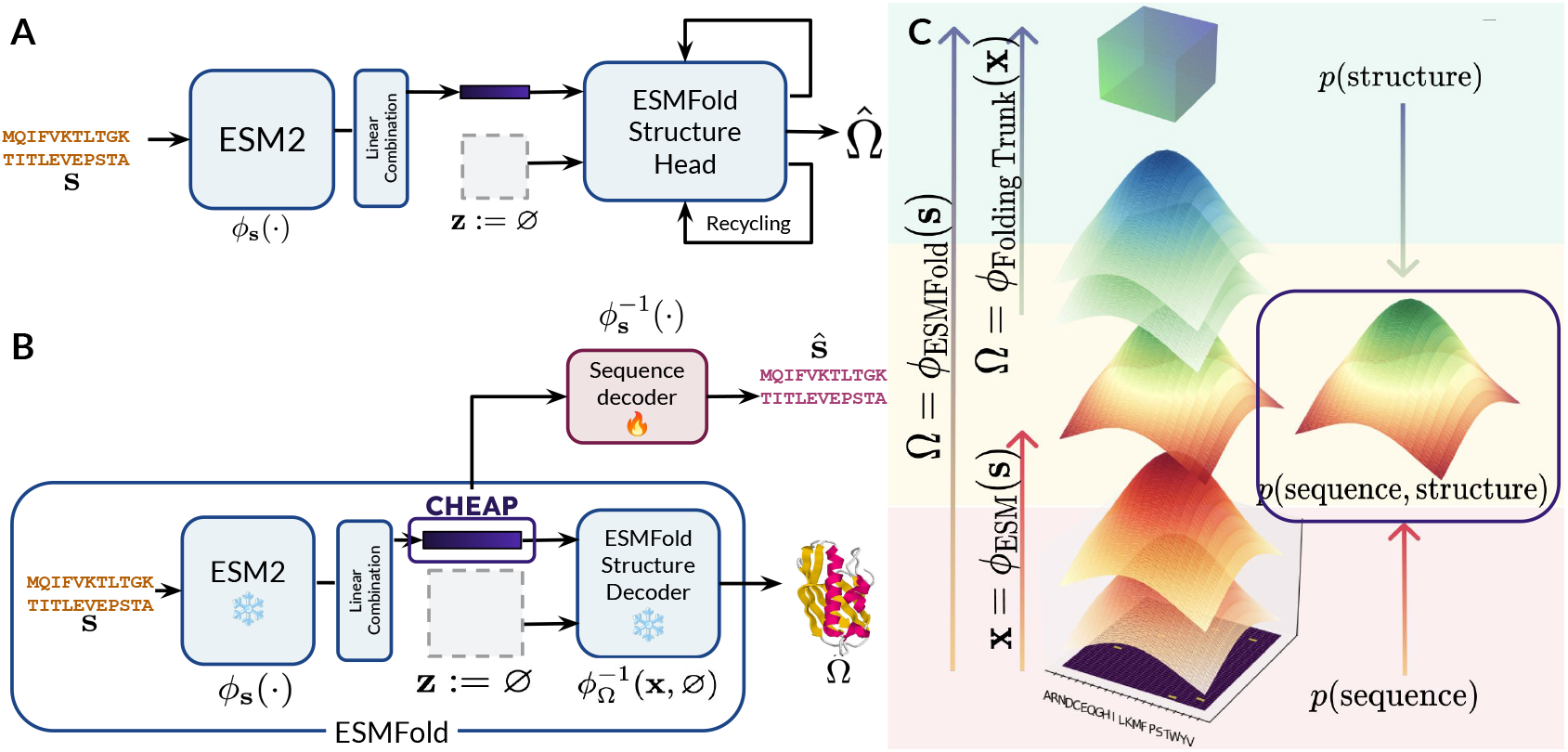
Deriving a joint latent space of structure and sequence using single sequence sequence-to-structure predictors. **(A)** Inference time usage of ESMFold, reinterpreted from Lin et al. [24]. A protein language model *ϕ*_**s**_(**s**), and a linear combination of output representations from each layer together maps a sequence **s** of length *L* to a latent embedding **x** *∈ R*^*L×*1024^. The Structure Trunk uses this input to predict an output structure, 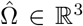, along with a *L × L* pairwise embedding, **z**, as input. At inference time, **z** is simply initialized to zeros, and updated in subsequent recycling iterations whereby outputs of the structure module, **s**^*′*^ and **z**^*′*^, are reused as inputs. **(B)** Harnessing the ESMFold latent space to obtain a joint representation of all-atom structure and sequence from sequence input. To decode the latent as structure (e.g. for latent generation), the original ESMFold structure decoder can be used; to map back to sequence, we separately train a sequence decoder, which achieves 99.7% validation accuracy. **(C)** The latent space **x** can be viewed as a joint embedding of sequence **s** and structure Ω; since the embedding requires only sequence to derive, but can be directly mapped to structure with the frozen ESMFold Folding Trunk, the embedding encodes both sequence and structural information.

Naïvely intercepting this latent space, however, presents numerous challenges. We find that pLM latent spaces contain abnormally high activations in certain channels that persist regardless of the input sequence (Figure 2), rendering them unwieldy for many downstream tasks. Additionally, the intrinsic dimensionality of language models, including protein language models, is often much smaller than the actual channel dimension [40]. Though subtle, embeddings with unnecessarily large dimensions sharply limits their utility. Aside from increasing computational resource demand, they limit the range of possible downstream applications. For example, high resolution visual synthesis involving larger pixel arrays is generally a more difficult generation task. For protein similarity quantification and search, the dot-product matrix multiplication performed between sequence embeddings has memory constraints that scale in 𝒪(*n*^2^) with the number of channels [3]. Furthermore, existing protein representations typically share the same length *L* as the original protein [24, 9]; however, as Transformer [43] architectures are being adopted for a broader range of tasks, it may be computationally desirable to downsample along the length dimension, since attention requires 𝒪(*n*^2^) memory with length *L*. Finally, the compressibility of data provides insight into its complexity and can inform downstream modeling choices; for example, the compressibility of data has been related to the compute-optimal frontier between favoring dataset size as opposed to scaling laws [29].

**Figure 2:**
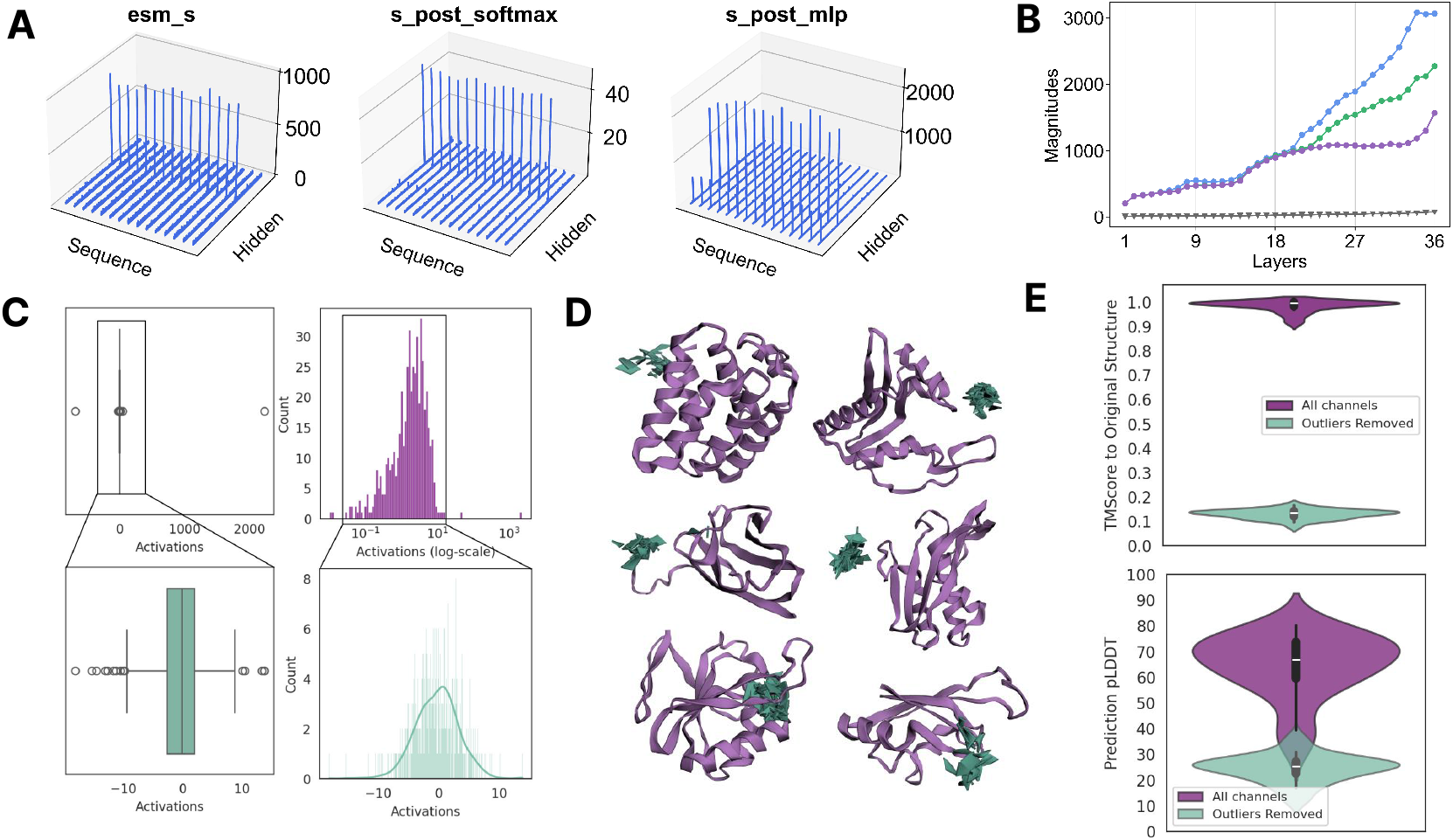
ESM2 contains massive activations in its embeddings. **(A)** For certain activations, massive activations occur at certain channels regardless of the sequence positions, at channel positions [274, 641, 37]. The pattern persists across layers (Appendix A). **(B)** Layerwise analysis demonstrating an increase in the magnitude of activations in the 3 billion parameter ESM2 model. Gray denotes the median activation magnitude; blue, green, and purple denote the top 3 largest activation magnitudes per layer output, and are significantly larger than the median activation. **(C)** Histogram of per-channel means. The total range of embedding values is greatly enlarged by the presence of massive activations. By removing three outlier channels with mean absolute values >20, the resulting latent space is closer to the Gaussian assumption necessary for diffusion. **(D)** Structure prediction results after setting the three outlier channels to zero. Purple denotes original ESMFold prediction, and green denotes ESMFold prediction after setting outlier channels to zero. Removing 3 channels with massive activations causes performance to deteriorate. **(E)** Quantitative assessment of the effect of outlier channel removal on structure prediction, measured by the TMScore (measure of structure prediction accuracy) and pLDDT (measure of model confidence), visualized for a random sample of 64 CATH structures. Model performance deteriorates by both metrics.

In this work, we compress the ESMFold latent space and introduce **CHEAP** (<u>C</u>ompressed <u>H</u>ourglass <u>E</u>mbedding <u>A</u>daptations of <u>P</u>roteins) representations in two variants: (1) token representations of the discrete biophysical concepts which induces the continuous 3D protein structure, and (2) continuous compressed embeddings that investigates the intrinsic dimensionality for tasks of interest. Continuous CHEAP embeddings can retain structural information at Angstrom-scale accuracy and achieve near 100% accuracy on retaining sequence information, despite reducing the channel dimension by up to 128×, and reducing the length of the embedding (i.e. downsampling) by 8×. Discretized CHEAP tokens also retain high levels of structural information, but unlike concurrent work such as ESM3 [13], CHEAP embeddings can be *obtained from sequence alone*, thus greatly enlarging the size of usable training data for applications requiring structural information. Our investigation furthermore sheds light on the mechanistic interpretability of increasingly popular protein language models, and on the information content asymmetry between structure and sequence, by demonstrating that structural information is much harder to compress than sequence information in protein embeddings. Code will be made publicly available at https://github.com/amyxlu/cheap-proteins.

## 2. Results

### 2.1. Defining a Joint Structure-Sequence Latent Space

A protein is comprised of a combination of 20 amino acids. Given a protein of length *L* with one-hot encoded sequence **s** ∈ 𝕀^*L×*20^ and 3D structure Ω ∈ ℝ^3^, our goal is to find an embedding that encapsulates the joint distribution **x** ∼ *p*(**s**, Ω); that is, the embedding **x** should both encode sequence information **x** = *ϕ*_**s**_(**s**) and structure information **x** = *ϕ*_Ω_(Ω).

We first decompose the mapping as

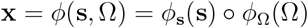

Though we can train separate encoders for *ϕ*_**s**_(·) and *ϕ*_Ω_(·), it is advantageous to leverage large pretrained models to improve performance and reduce computational overhead. In particular, ESMFold demonstrates that structure prediction is possible from protein language model representations learned by sequence masked language modeling, which reduces computation requirements and improves performance for orphan proteins [24]; we use this model as a representative for sequence-to-structure models. The ESM2-3B language model *ϕ*_ESM_(**s**) embeds sequence **s** into a 2560-dimensional sequence representation, which then is mapped via linear projections to an embedding **x** *∈* ℝ^*L×*1024^. This is used as input to the Folding Trunk 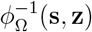, where 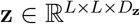 represents pairwise contacts. An overview of the original ESMFold model with notations used in this work is shown in Figure 1A.

A key observation is that at inference time, it is empirically sufficient to initialize input **z** as an array of zeros to obtain high quality structure predictions as reported in Lin et al. [24], such that 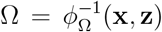 becomes 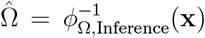. That is to say, **x** is a direct inverse-mapping from 3D structure Ω to a high-dimensional latent space. This sequence embedding **x** therefore, obtained as the latent space of the ESMFold model, can be seen as our embedding of the joint distribution of *ϕ*(Ω, **s**):

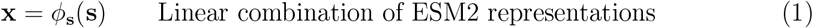

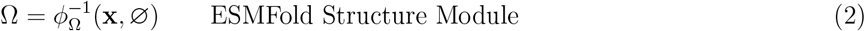

Note that **x** is not precisely the same as ESM2 embeddings, since in ESMFold, ESM2 representations from each layer are linearly combined with trainable weights to create the **x** embedding that is input to Folding Trunk 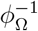. Appendix A provides a code sketch for obtaining **x** ∼ *p*(**s**, Ω), and a graphical description is shown in Figure 1B.

We would also like to map this latent space back to sequence and structure space, for assessing compression performance and for generative modeling. To decode this latent space back to protein sequences, a sequence encoder 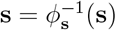 is trained separately. We achieve a per-token accuracy of 99.7% on a randomly partitioned heldout set (detailed in Section 4.5), likely because the task of mapping a language model embedding to 21 classes is fairly simple, and closely related to how the ESM2 model was trained. To obtain structure from embedding, we can simply use the frozen ESMFold folding trunk, 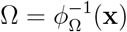. If generative model parameters *θ* are learned to approximate *p*_*θ*_(**x**), then at inference time, after sampling 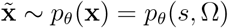, we can generate new 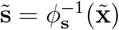 and 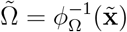, using the decoders 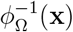 and 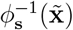 defined above.

### 2.2. ESMFold Latent Space Contains Massive Activations

We find that the latent space of ESMFold is pathologically disorganized, whereby some channels have mean values up to 3000× larger than others. This is consistent with reports of this phenomenon in large Transformer [43] models for text and image modalities, where they are sometimes referred to as massive activations [37] or outlier features [7]. Figure 2A visualizes embedding values at various intermediate layers between the ESM2 output and the folding trunk, following variable notations in Appendix A.

Massive activation accumulation begins in the ESM2 language model. The top values in any given layer can be up to 3000× larger than the median activation value in a given layer (Figure 2B). Figure 2C visualizes the means of each channel. When three outlier channels with massive activations are removed, the resulting distribution is more organized; however, removing these three features alone causes structure prediction performance to deteriorate, as seen qualitatively (Figure 2D) and quantitatively (Figure 2E), bringing the TM-Score of predictions down from an average of 0.97 down to 0.14.

Massive activations limits the flexibility with which large protein models can be used as foundation models. Dettmers et al. [7] examines 8-bit model quantization to reduce memory usage during inference, which is rendered difficult due to the expanded numerical range caused by the 0.01% of abnormally high values. Massive activations have also been observed to dominate attention patterns, and have been suggested to act as an implicit bias term [37]. Given that attention mechanisms have been shown to learn protein contacts [30], these massive activations may play interpretable roles in how structure information emerges from protein language models; we leave the specifics of this investigation as future work. We combat this by applying a per-channel normalization scheme, described in Section 4.1.

### 2.3. A Neural Compression Architecture for Protein Embeddings

The equivalence between data compression and data distribution representation learning is foundational to machine learning [25]. Biological data is comprised of both signal and noise, not all of which is necessary to understand its structure or function; despite this, data compression remains understudied for proteins. Our work aims to bridge this gap by studying protein embeddings as an instance of neural compression. We design the Hourglass Protein Compression Transformer (HPCT) architecture, an autoencoder with a bottleneck layer, further detailed in Section 4.2 and visualized in Figure 3, for protein embedding compression.

**Figure 3:**
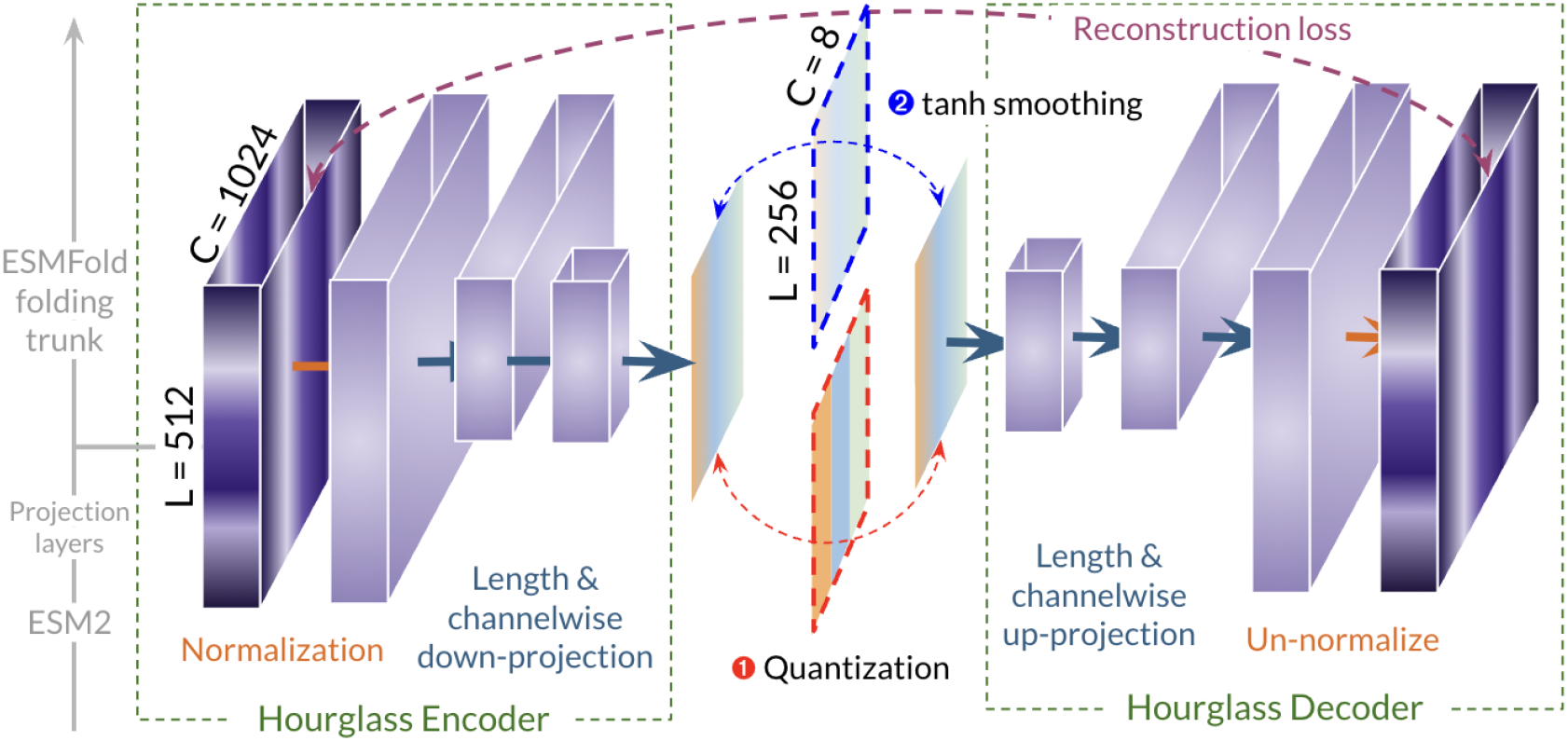
The Hourglass Protein Compression Transformer (HPCT) model. For a protein with 512 amino acids, the embedding right before the ESMFold structure encoder has dimensions 512 × 1024. To correct for massive activations, the embedding is normalized using the statistics of each channel (Section 4.1). The encoder condenses the channel dimensions with a linear projection and a linear downsampling operation along the length dimension. In the bottleneck layer, we examine methods for obtaining both discrete and continuous compressed embeddings.

Proteins have different lengths, rendering the ResNet [14] autoencoder architecture used in vision a poor choice. Inspired by Nawrot et al. [28], HPCT includes a linear downsampling operation (Algorithm 8) which “shortens” the embedding *x* ∈ ℝ^*L×D*^ to 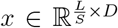 using a shortening factor of *S*. A linear projection further compresses the channel dimension. The model is trained end-to-end with a mean-squared error reconstruction loss, such that it is trained to maximally preserve the original content across the bottleneck layer. At inference time, the output of the bottleneck layer is used as the compressed embedding. In early experiments, we also apply a sequence-space loss (i.e. cross-entropy loss constraint that decoded amino acids corresponds to the original) and structure-space loss (backbone Frame-Aligned Point Error [20]), but did not find these to substantially improve performance.

#### Continuous vs. Discrete Compressions

Performing data compression also has practical motivations. For example, in high-resolution image synthesis, increased number of pixels exacerbates FLOPS needed and generation difficulty. To combat this, Latent Diffusion Models (LDM) [32] performs diffusion modeling [17, 36] in a continuous compressed latent space learned by a VQ-GAN [10] encoder. Another successful technique is encoding continuous images as discrete tokens, and generate tokens using an autoregressive model or by sampling from mask tokens [5, 15, 31]. We aim to make contact with these paradigms by investigating how to build the necessary autoencoder for protein data.

All experiments uses the embedding **x** defined in Section 2.1, though the architecture can be used for any sort of embedding compression. The dimensions of **x** defined in Section 2.1 is *L* × 1024 for a sequence with length *L*.^1^ For a protein sequence of length 512, the size of this embedding maps to high-resolution image synthesis. As a point of comparison, LDMs [32] compresses input images of size 256 × 256 × 3 down to 64 × 64 × 4 before diffusing.

At the bottleneck layer of HPCT, we prefer either quantization step at the bottleneck to obtain **discrete** compressed embeddings, or a tanh step to obtain **continuous** compressed embeddings. The tanh layer constrict embedding values to between [−1, 1] and improve its utility (e.g. for latent diffusion); in early experiments, we found that it does not degrade performance.

#### Compression Evaluation Metrics

In following sections, we assess continuous and discrete compression performance. To assess in-distribution structure reconstruction, we train and assess on the CATH [35] database, though a sequence-only dataset such as Pfam [27] or UniRef [38] can also be used (select results in Appendix C). Mirroring the usage of perceptual losses for assessing compression in images [10], which favors retention of visual details necessary for downstream tasks as opposed to purely reconstruction, we examine reconstruction performance in the structure and sequence space in addition to the mean-squared-error between input and reconstruction. The template modeling score (TM-Score) is a backbone-only metric of structure reconstruction, while root-mean-squared distance (RMSD) is a more fine-grained measure between atom positions. Root-mean-squared paired distance (RMSPD) is similar to RMSD, but uses pairwise distances rather than atom positions; we use it to assess superimposition-free reconstruction accuracy. Sequence reconstruction accuracy is the fraction of token matches after decoding back to sequence space, divided by the number of tokens in the sequence. As reference points for interpreting structure reconstruction results, we note that 1.34Å is the inter-carbon distances taken from Jumper et al. [20], and range of 0.8Å to 4.1Å is the range experimental resolution for the original structures.

### 2.4. A Discrete Vocabulary of Protein Structure and Function

“Quantization” or “tokenization” here refers learning an encoder that maps an input to a discrete representation. Given a *d*-dimensional input **a** ∈ ℝ^*d*^, and a pre-specified codebook size *C*, the goal is to find a set of integers from 𝒞 ∈ {0, 1, …, *C*} that represents the input. Intuitively, though pixel inputs of the visual world is continuous, many discrete concepts exist, such as color, shape, and size; similarly, though structure exist in the continuous 3D space, the biophysical concepts that cause the shape might manifest discretely. Furthermore, obtaining a discrete vocabulary of representations allows scaling infrastructure for large language models to be applied.

Tokenized representations of protein structure is an area of limited investigation. Fold-seek [42] introduces the 3Di vocabulary for its search algorithm, though this is limited to 20 amino acids, and it is known that codebook size (i.e. number of discrete options) is a crucial factor in representation quality. Some concurrent works have also examined characterizing a tokenization scheme for structure [11, 23, 12]. However, these methods are learned on structural input, unlike the scheme here, which can be used on sequence-only input. ESM3 [13] constructs separate discrete vocabularies for structure, function, and sequence, as opposed to directly crafting a single discrete embedding that can be decoded into multiple modalities.

While Vector-Quantized Variational Auto-Encoders (VQ-VAE) [41] is a popular choice, it can be difficult to optimize, and is prone to “codebook collapse”, whereby a few codes are over-utilized, and require specific intervention [39, 21, 8, 19]. Thus, we also investigate the recently proposed FSQ [26] approach, which obtains token representations without quantizer parameters. Details of the VQ-VAE and FSQ methods are described in Section 4.4.

The choice of codebook size relates to the bits of information content in the data, and the degree to which information loss is tolerable for the downstream task. It is unclear how to best select this for all-atom protein structure and sequence. We find that while VQ-VAE outperforms FSQ on codebook sizes smaller than 2^8^ = 256, FSQ consistently outperforms VQ-VAE on latent space reconstruction MSE, structure space metrics (both the backbone-only TM-Score and the all-atom RMSD), and sequence space metrics (Figure 4). This is consistent with image experiments in Mentzer et al. [26]. FSQ generally has more evenly distributed codebook usage, especially for larger codebooks. This is reflected in the high VQ-perplexity (perplexity when selecting a code during the VQ-VAE quantization process) as codebook size *C* increases, a metric which does not apply to FSQ.

**Figure 4:**
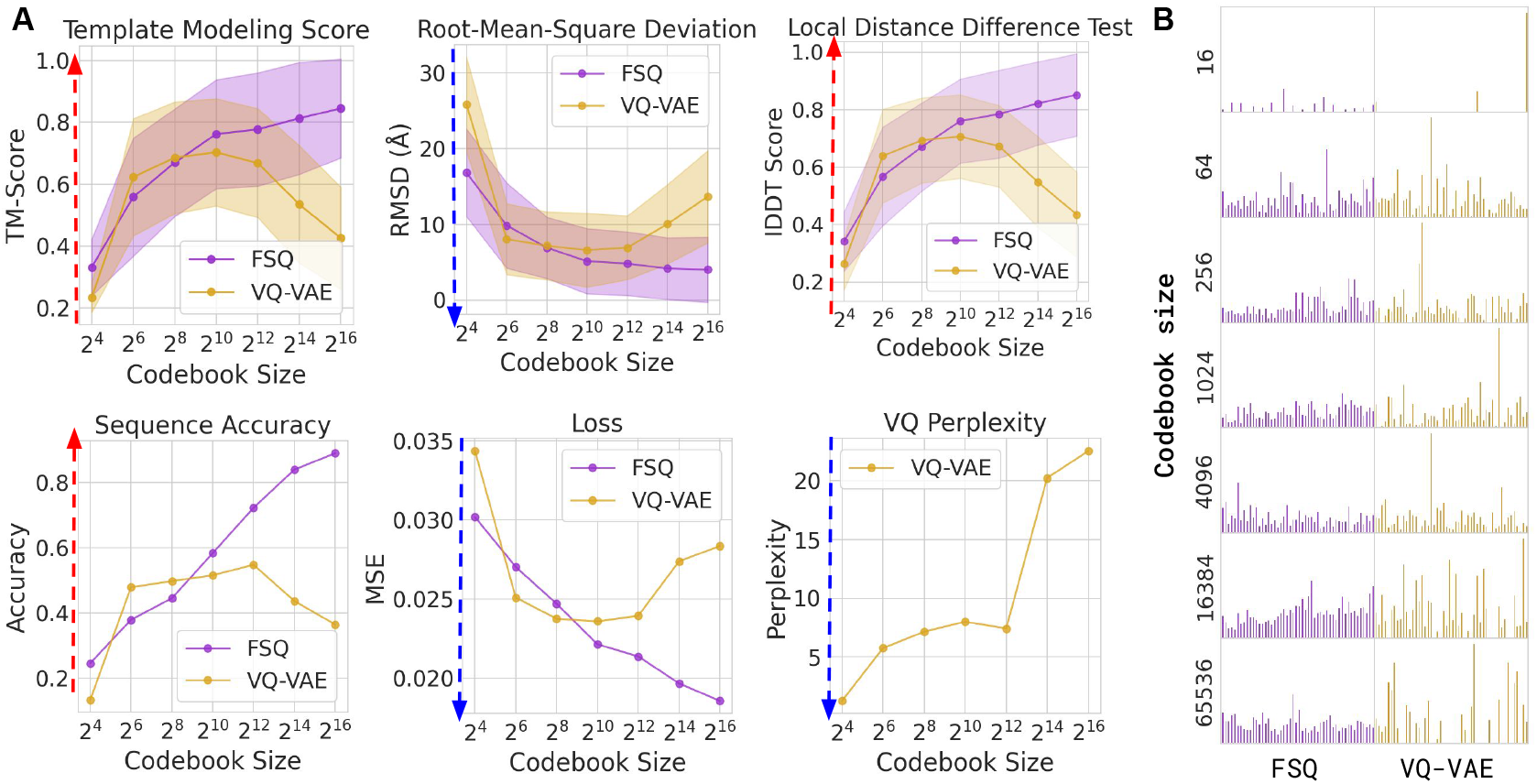
Assessing structure, sequence, and latent reconstruction across codebook sizes for the FSQ and VQ-VAE methods. **(A)** Across all metrics, FSQ outperforms VQ-VAE at codebook sizes greater than 2^10^. Blue arrows denote metrics where lower is better, and red arrows denote metrics where higher is better. VQ-perplexity refers to the codebook selection prediction in VQ-VAE, which does not apply to FSQ. **(B)** Examining codebook utilization for FSQ and VQVAE. We find that codebook utilization is generally more favorable using FSQ (left), though we do not find catastrophic codebook collapse for VQ-VAE above a codebook size of 2^4^.

### 2.5. Continuous Compression Performance on Structure Reconstruction and Function Prediction

Our experiments empirically validate the hypothesis that many channel dimensions in protein representations are extraneous and that the intrinsic dimensionality likely lower. Figure 5A examines the ability for compressed embeddings to retain structural information, examined both with and without length downsampling. Even with a 32× compression factor down to only 32 channels, the model is still able to achieve <1.34Å RMSD in structure fidelity compared to the original prediction. When shortening by a factor of two, one can attain a <1.34Å RMSD at 64 channels or higher. For sequences, there is nearly no drop in reconstruction performance until compressing to less than 8 channels. Interestingly, the largest drops in performance is found after compressing to 4 channels, by both structure and sequence reconstruction accuracy, which may be related to the average number of atoms in amino acid residues. The pairwise RMSD, which is independent of superimposition, is generally lower than the RMSD, suggesting that some atomic distances may be due to loss of precision on trivial 3D super-imposition, rather than insufficiently learning protein information. A visual depiction of how compression changes the predicted structure is shown in Figure 5C. Appendix E shows TSNE results for embeddings at different levels of compression.

**Figure 5:**
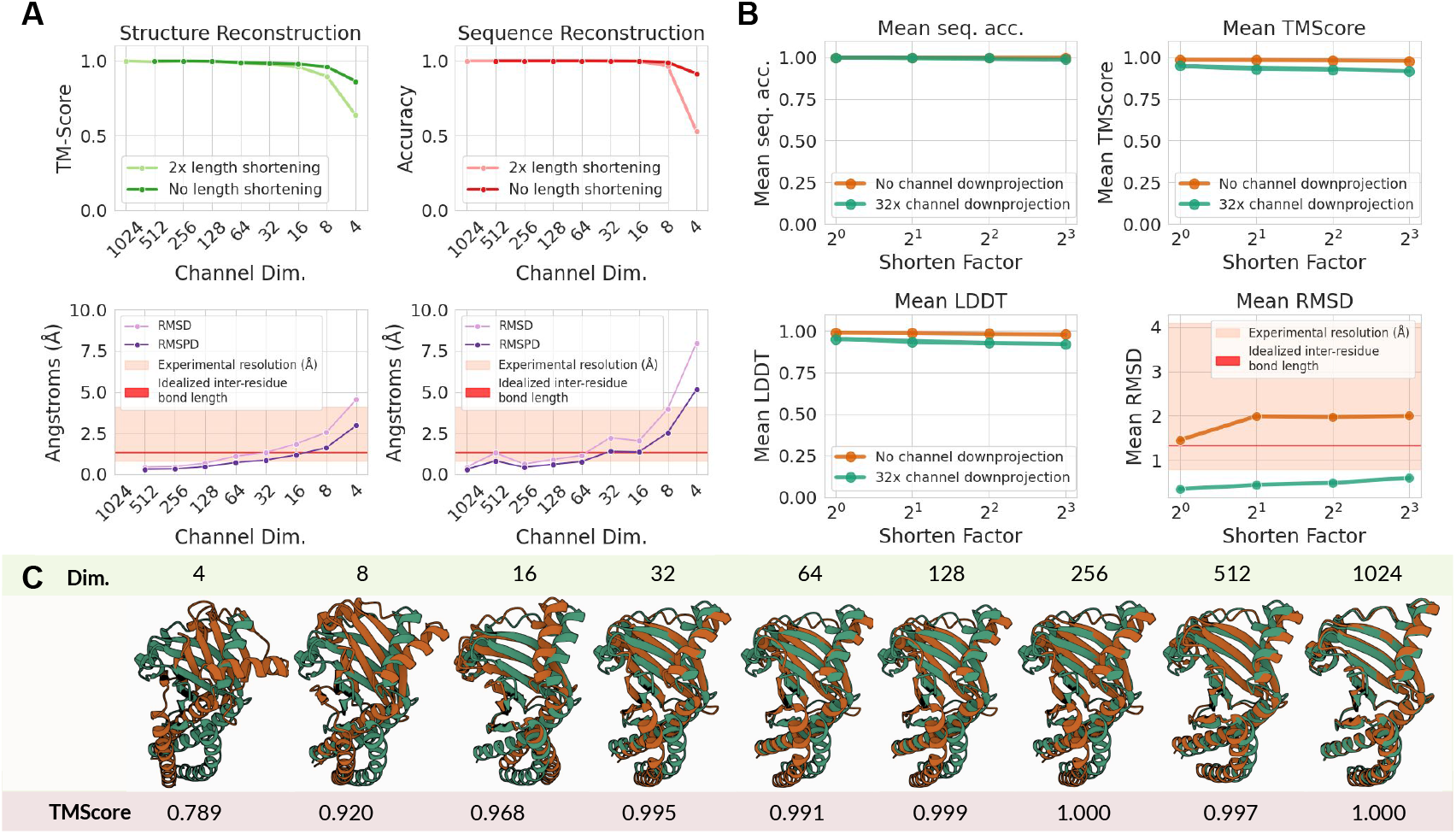
Assessing compressibility of structural and sequence information in protein embeddings. **(A)** Examining compressibility of backbone structure (TM-Score), sequence identity (sequence recovery accuracy), all-atom structure (RMSD), and superimposition-free all-atom structure (pairwise RMSD). Sequence information and backbone information can be preserved nearly perfectly even after reducing down to only 8 channels; all-atom structure RMSD similarly achieves <1.34Å accuracy at 16 channels or less. **(B)** Length information can be compressed while preserving sequence and backbone information even after reducing channel dimensions to 32. **(C)** Visual overlay of structural information compressibility. Even with only 4 embedding channels, secondary structures can be preserved after reconstruction.

We further examine function and localization prediction using datasets from the PEER benchmark [45] for embeddings at different compression levels, and fund that functional information seems to suffer more with compression than structure and sequence information. Section 4.5 describes the details of the training procedure. This is consistent with recent works demonstrating that protein language models fail on downstream tasks which do not involve structure [22]. For some tasks, such as *β*-lactamase activity prediction and subcellular localization, compression seems to be more impactful than tasks such as solubility. We show comparisons against other methods along with additional benchmarks in Appendix D. On structure related tasks, compressed embeddings can compare competitively with other methods, sometimes exceeding their performance.

### 2.6. Interpolation and Noising in the Latent Space

To examine the smoothness of the latent space, we perform linear interpolations in the embedding space, and examine how this manifests in the structure and sequence space. For two real proteins in the CATH dataset, we obtain the CHEAP embedding representation, and perform a linear interpolation as **x**^*′*^ = *t***x**_1_ + (1 − *t*)**x**_2_. We then transform this embedding back to the sequence and structure space, and examine both their similarities to the proteins which they were interpolating between, and measures of the “naturalness” of the produced sequence via perplexity and of the structure via pLDDT (Section 4.3).

Figure 7A demonstrates reconstruction results in the latent space. We find that the unprocessed ESMFold latent space is not very “smooth” – that is, despite changes linearly in continuous space, the discrete changes in the sequence space is limited, as visualized by the sequence accuracy to the end sequence. A similar case is observed for structures. After the per-channel normalization (Section 4.1) and compressing down to 8 channels, the sequence space is more “smooth” and changes more gradually with linear interpolation in the embedding space; however, the structure space remains less smoothed out. Despite being a smoother interpolation between the two natural proteins, these proteins are less “natural”, though it may better reflect the ragged nature of the hypothesized protein fitness landscape [33]. This may reflect the classic quality-versus-diversity trade-off, but also raises questions on the geometry of the protein structure fitness landscape, and whether a more rugged or sharp landscape is more suitable for exploration. Figure 7C provides a visual of how the structures change when interpolating in the original latent space versus the compressed latent space. The jumps are more gradual when interpolating in the compressed latent space and then transforming back to the embedding with massive activations; however, they also tend to be more unstructured.

**Figure 6:**
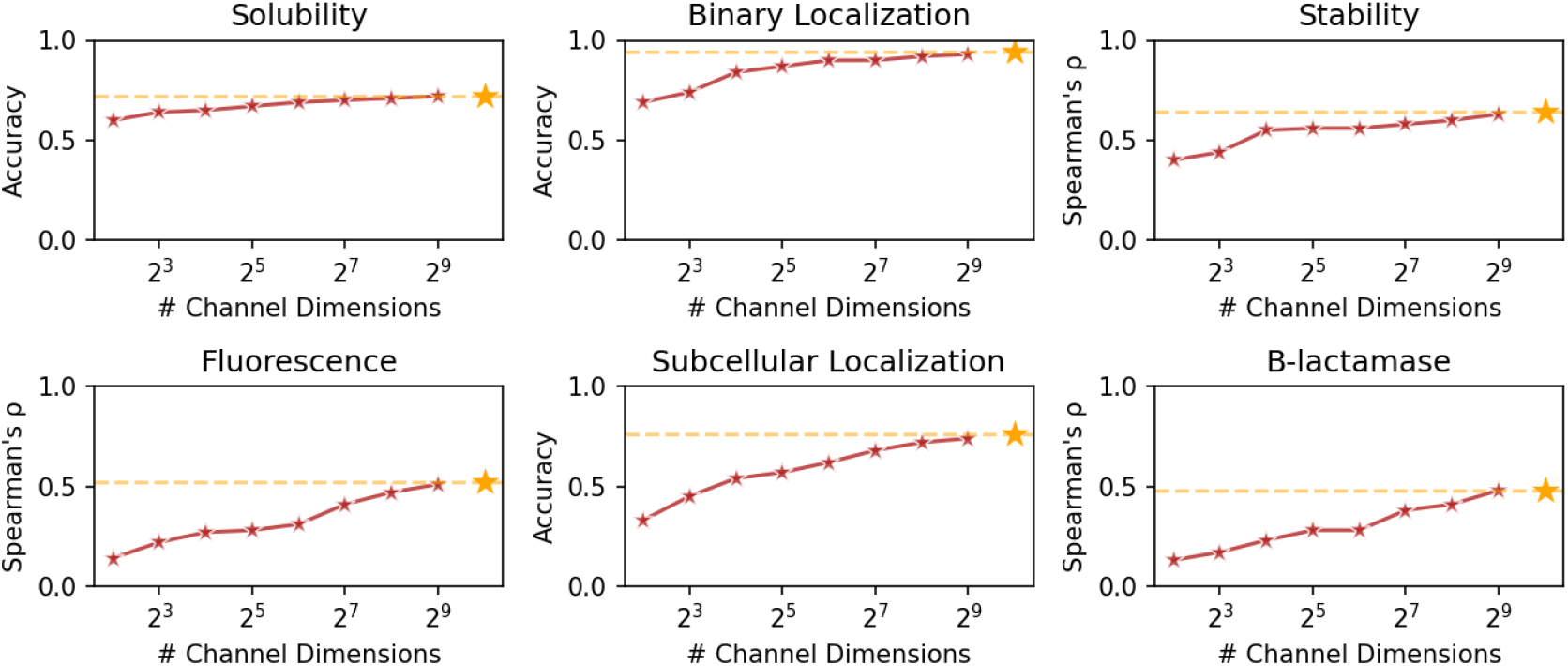
Using linear probes to evaluate compressibiliy of functional information, with 2× lengthwise shortening applied. Orange star denotes uncompressed performance. Unlike structure and sequence, performance degrades more gradually for some functions (e.g. *β*-lactamase activity), though the performance drop is less pronounced for other tasks (e.g. solubility).

**Figure 7:**
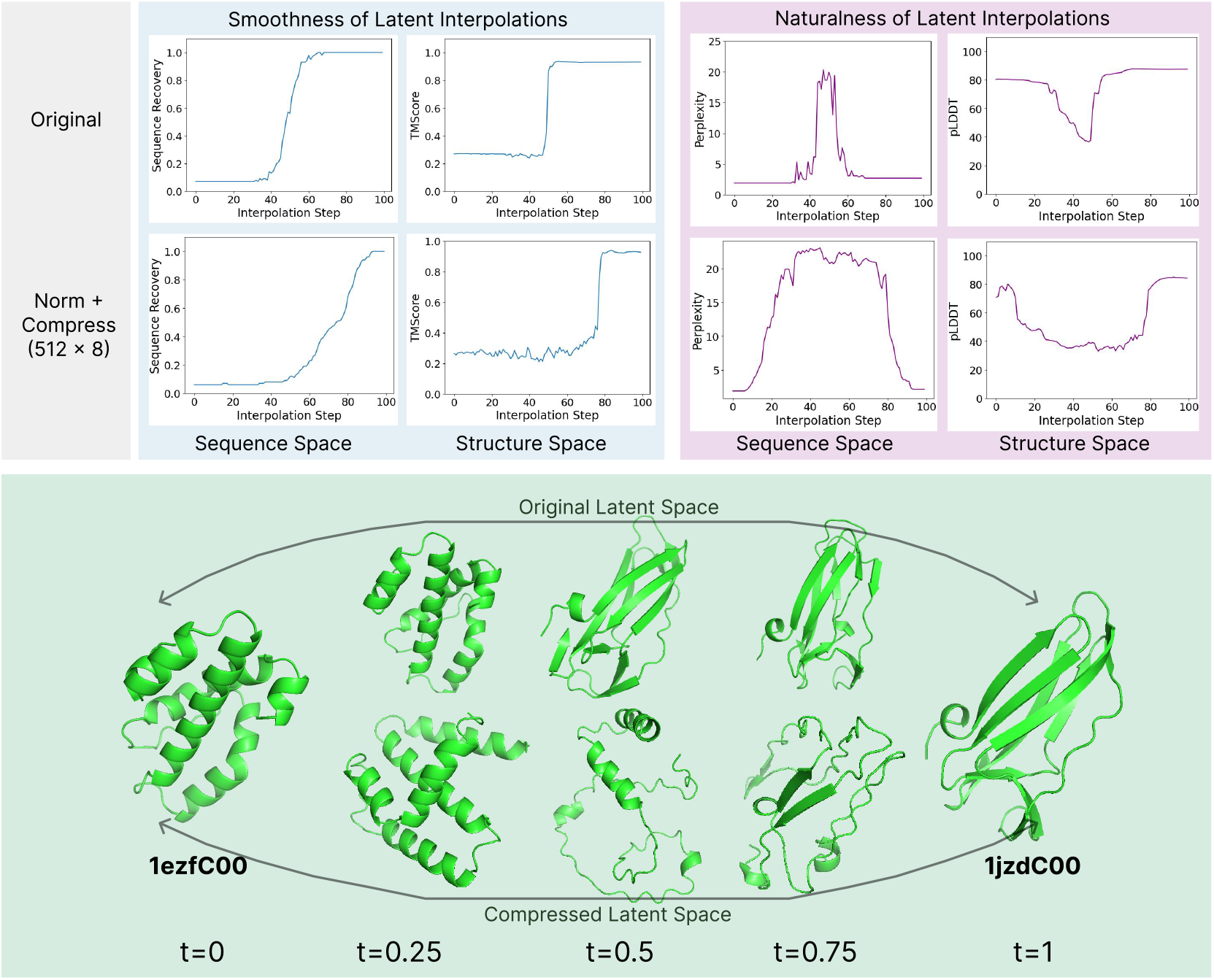
Comparing linear interpolation in various embedding spaces as empirical assessment of geometry. **(Top)** To examine how smoothly sequence changes as a result of interpolating in a given latent space, we decode interpolations back to sequence/structure, and examine sequence recovery accuracy (↓) and backbone TMScore (↑) to the final sample. Despite moving linearly in embedding space, the actual sequence and structure samples that embedding interpolations map to remain tightly coupled to proteins seen in the training set. Interpolations typically occupy regions of low perplexity (↓) and high structure prediction confidence (pLDDT, ↑). Our compression and normalization schemes hele improve the smoothness of interpolations and raggedness of the approximated fitness landscape. **(Bottom)** Visual examination of structure reconstructions of linear embedding interpolations.

## 3. Discussion

In this work, we characterize how the latent space of large protein sequence-to-structure models can represent the joint distribution of sequence and structure. We further identify pathologies of massive activations in ESMFold; indeed, the original paper uses a linear combination of representations from all layer outputs of the language model rather than the final embedding, and though no explicit justification or ablations were given in Lin et al. [24], our findings of how pathological massive activations accrue across layers may contribute to why representations from intermediate layers might be as or more informative than the final layer embedding. We also find that protein sequence embeddings are highly compressible, and develop the Transformer-based HPCT autoencoder architecture. Sequence information can be captured by embeddings of only a 8 channels. All-atom structural information is less compressible than sequence information, but nonetheless, remarkable reconstruction performance can be obtained by embeddings that undergo 128× channelwise compression and 8× lengthwise compression between amino acids. We also develop a *structure tokenization dictionary* which can be *obtained from sequence alone*, unlike concurrently proposed structural tokenizers [11, 23, 12]. Further, we show that FSQ is more effective than the dominant VQ-VAE approach (which has also been used for proteins by recent works [13]), especially for large codebooks.

Building a representation of the joint distribution of structure and sequence has many desirable traits for generation, search, and representation learning. The CHEAP series of embeddings is a compact joint embedding of the latent space that can be flexibly employed. Furthermore, our empirical results raise interesting theoretical questions on the information content and geometry of protein embeddings. The compressibility of embeddings with respect to sequence identity and structural precision corroborates empirical investigations that the intrinsic dimensionality of existing structure foundation models is likely much lower [40], and there may be over-parameterization in existing models. The lack of a clear pattern for linear probes relating to function further illustrates the poorly understood nature of how protein embeddings capture relevant information.

An alternative means of achieving this is to train and/or finetune ESMFold (or another sequence-to-structure model) end-to-end with the bottleneck layer and with a channel normalization before it. We hypothesize that this should improve the properties desirable for empirical downstream use without compromising structure prediction quality, though due to computational resource limitations, we leave this as future work.

## 4. Methods

### 4.1. Per-Channel Normalization

To address the issue of massive activations as shown in Figure 2, we use a per-channel normalization scheme. Conventionally, following Ho et al. [17], image generation works rescale image pixel values from [0, 256] to [−1, 1]. Subsequent works find this bounding strategy to be important to performance, such as color saturation [34]. To remedy channel-specific massive activations, the embeddings are processed as:

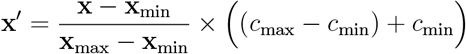

where **x**_min_ and **x**_max_ are vectors with shape (1024,), and broadcasted along length dimension. They denote statistics calculated independently for each channel. This prevents outlier channel values from dominating the normalization. In experiments, we choose *c*_min_ = *−*1 and *c*_max_ = 1.

Visual inspection shows that after this transformation, the distribution better befits the Gaussian distribution needed for diffusion, as visualized in Figure 2. A random subset of 5,000 samples from CATH [35] is used, but we find that these mean and standard deviation statistics vary little across datasets such as CATH, UniRef [38], and Pfam [27], consistent with the general observation that massive activations are not input-specific [37]. In simple diffusion experiments, this greatly improves stability of training.

### 4.2. Hourglass Compression Transformer

A sketch of the hourglass compression transformer is as follows, inspired largely from Nawrot et al. [28]. Linear downsampling layers serve to shorten the embedding lengthwise, while the downprojection reduces the channel dimension. The attention resampling is an attention layer that attends to both the pre- and post-shortening embeddings. This is similar to the approach in LDM where compressed latent space is taken to be the continuous representation just prior to the VQ layer (except we replace the VQ with FSQ in this case).

#### Algorithm 1

Hourglass Compression Transformer

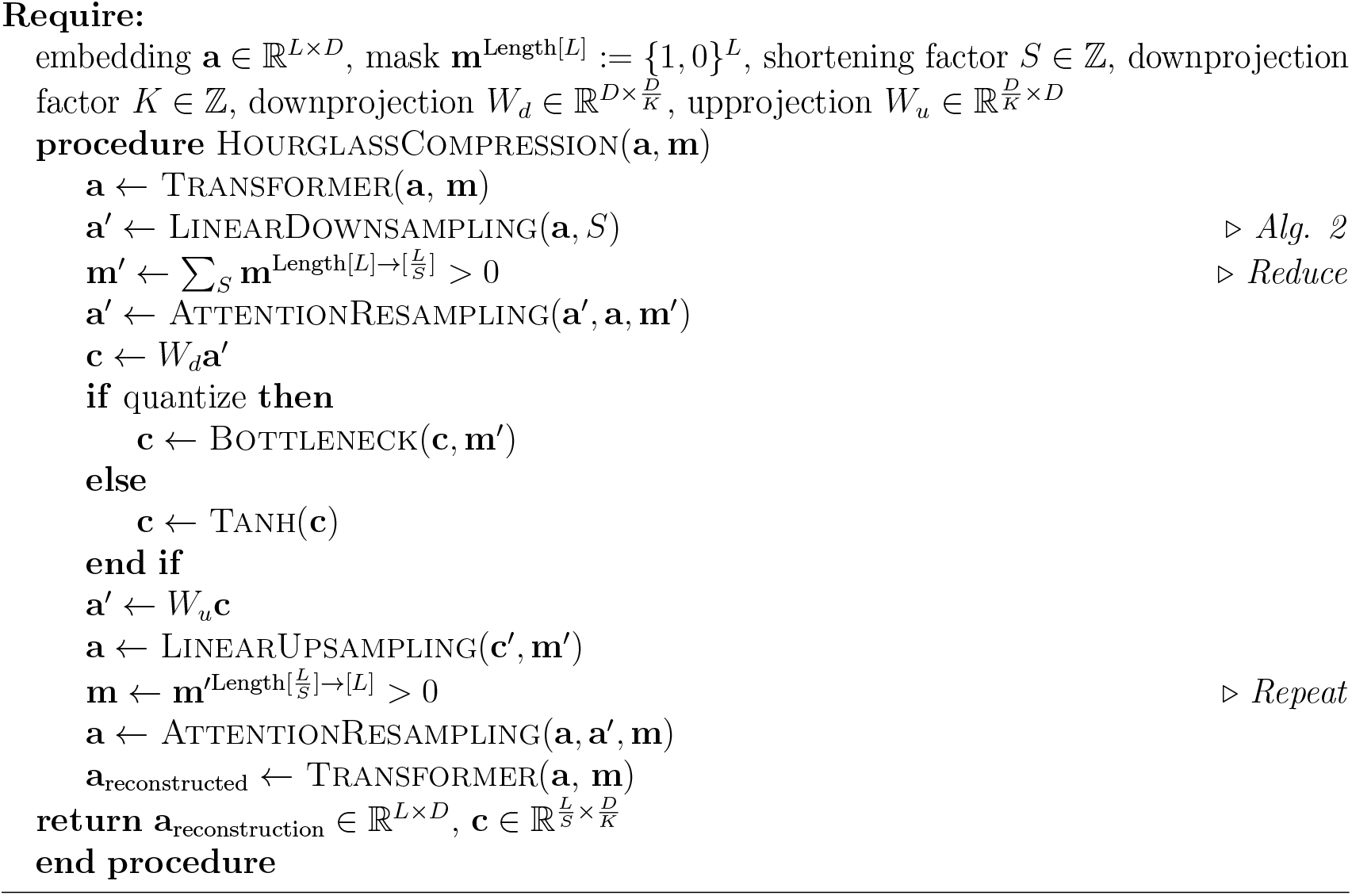

Though different operations may be used for *g*(*x*), we find a simple linear downsampling to work well (Algorithm 8), which can be seen as a convolution with a filter size and stride both equal to *S*. Since the original model is designed for sequence-to-sequence tasks rather than compression, we remove the skip connections that would make the solving the reconstruction task trivial. Additionally, we add a projection layer along the channel dimension after each shortening operation. At training time, **x**_reconstructed_ is used for calculating the mean-squared-error reconstruction loss, and at inference time, the output of the encoder is used as the compressed representation, with additional processing in the bottleneck, depending on if the compression is discrete or continuous. Linear downsampling consists of flattening operation, followed by a learned downsampling weight matrix, and finally reshaped back to *L/S*, where *L* is the original length, and *S* is the shortening factor.

#### Algorithm 2

Hourglass Encoder Shortening by Linear Downsampling

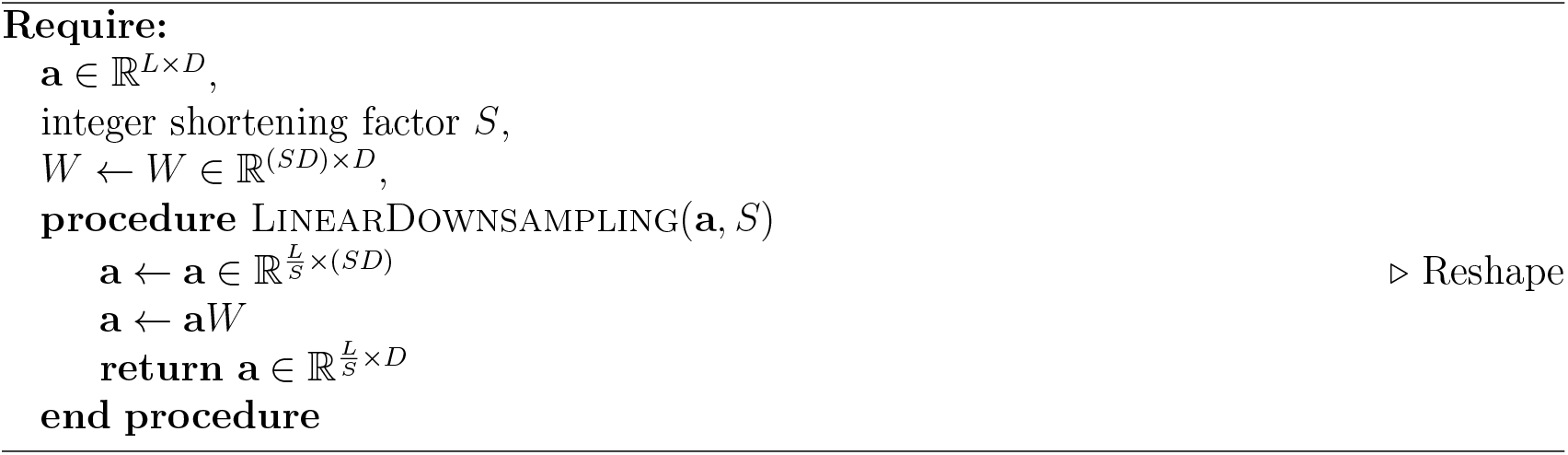

### 4.3. Naturalness Evaluation

For evaluating sequence naturalness with perplexity, we use the autoregressive RITA XL model [16]. Intuitively, perplexity can be seen as a measure of how “confused” a model is when selecting the next token. If a sequence is in-distribution with the natural proteins that the model was trained on, its perplexity will be lower, since the model can make sense of the patterns. Conversely, if a sequence out-of-distribution (e.g. is not a valid protein) and learned patterns from natural proteins cannot help the model discern what the next token should be, then the perplexity increases.

For evaluating structure naturalness, we use the predicted Local Distance Difference Test (pLDDT) score, which is returned by the ESMFold prediction head along with the all-atom structure. The metric was introduced in AlphaFold2 [20] and is a predicted measurement of the model’s certainty in the structure at that particular position. Similar to the reasoning behind perplexity as a metric of naturalness, a high pLDDT suggests that a metric is in-distribution with the natural proteins that was seen during sequence-to-structure training. Note that pLDDT tends to favor structured regions, and is typically low for disordered regions, even in real proteins.

### 4.4. Quantization Methods

#### Vector-Quantized Variational Autoencoder

The VQ-VAE [41] learns a discrete, compressed semantic representation of the input, typically of images. In the forward pass, the encoder *h*_*e*_ produces a continuous feature representation of input **x**. Then, each feature vector is mapped to a discrete code in the codebook space, 𝒞, where each discrete code is associated with a continous vector **e**_*i*_. The assembled array of learned codes and their vector embeddings **z** = {**e**_1_, **e**_2_, …**e**_|*𝒞*|_} and their corresponding feature features are fed into the decoder *h*_*q*_(**z**).

Since the quantization operation is not differentiable, the straight-through estimator (STE) [1] is used by copying the gradients from the decoder input to the encoder output. The codebook is selected via a nearest-neighbor search in Euclidean space; auxiliary losses are introduced to pull the code vectors towards the unquantized encoder outputs. As in autoencoder training, a reconstruction loss between output and input is also used. The complete VQ-VAE loss is:

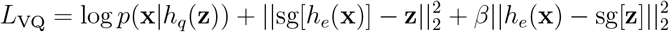

#### Finite Scalar Quantization

Rather than using a nearest-neighbor search to choose a code, FSQ directly quantizes the continuous encoder representations **z** *∈* ℝ^*d*^ into *L* bins:

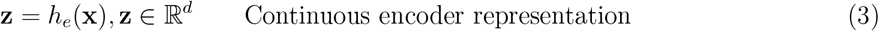

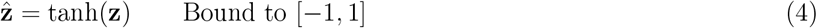

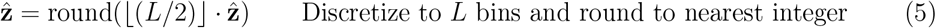

The predetermined bins *L* is selected to be small relative to VQ-VAE codebook sizes. The *implicit codebook size* |𝒞|, however, comes from the combinatorial possibilities arising from using one of *L* integers at each of the *D* channels. For **z** with *d* channels, there are *d* associated integer representations, and thus |𝒞| = *L*^*d*^. A large implicit codebook can thus be achieved, while forcing all codes to be used.

### 4.5. Training

#### Compression Model Training

For all continuous compression models, we use a learning rate of 8^*−*5^ with the AdamW optimizer and 5000 warmup steps. For discrete compression models, we use a learning rate of 1^*−*4^, with the AdamW optimizer and 1000 warmup steps. Train and test sets were split randomly. CATH is used for all experiments, with the exception of the experiments in Appendix C.

#### Sequence Decoder

For the sequence decoder, we use a fully connected network with two layers and a hidden size of 1024. This network is able to converge relatively fast. Though ESM2 was already trained with a masked token objective, and thus has pretrained weights that map the language model output to logit space, we opt for retraining, since our defined embedding is intercepted after the linear projection layers (with 1024 dimensions), where as the original ESM2-3B model has 2580 dimensions. The output has 21 classes (20 amino acids in addition to the unknown residue placeholder). In early experiments, we also try using an Transformer architecture for this decoder, but find that the over-parameterization actually decreases performance.

### 4.6. Function prediction

For results in Figure 5C and Appendix D, we use the dataset splits and tasks from Xu et al. [45]. For all compressed embeddings, we add a trainable projection layer back to 1024, such that the size of the downstream fully-connected network used for prediction is the same size for all compared networks. A mean pool is applied to the embeddings, with a 2-layer multilayer perceptron used as the downstream probe.

## 5. Acknowledgments

Authors thank Mingjie Sun for productive discussions on massive activations in large language models, and help with producing Figure 2A. Authors additionally thank Sidney Lisanza, Daniel Severo, Igor Mordatch, Andrew Leaver-Fay, and Lian Huang for discussion and comments. AXL is supported in part by the NSERC PGS-D scholarship and Nissan.

## Appendix A. Extracting Embedding from ESMFold

We provide a code sketch of how to obtain the embedding space used in this work, based on modified and annotated source code from ESMFold [24]. Note that `self.esm_s_combine` and `self.esm_s_mlp` are both trained end-to-end with loss objectives from the original ESMFold paper; these weights add more structural awareness to the embedding space which we examine in this work than the original ESM2 3B embedding.

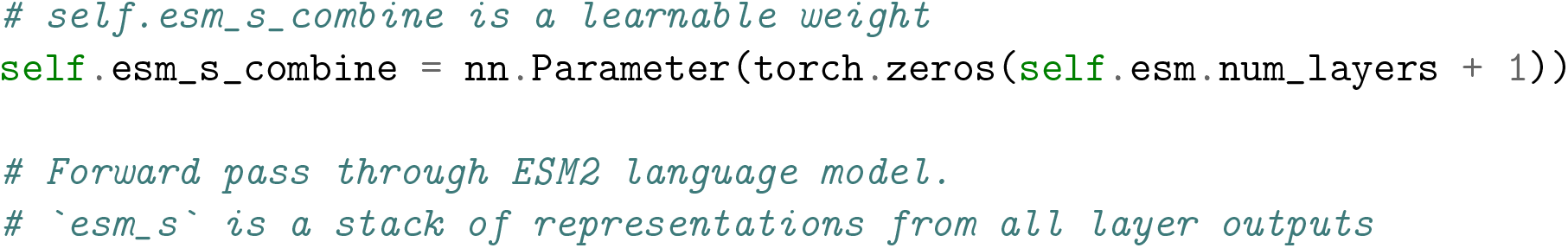
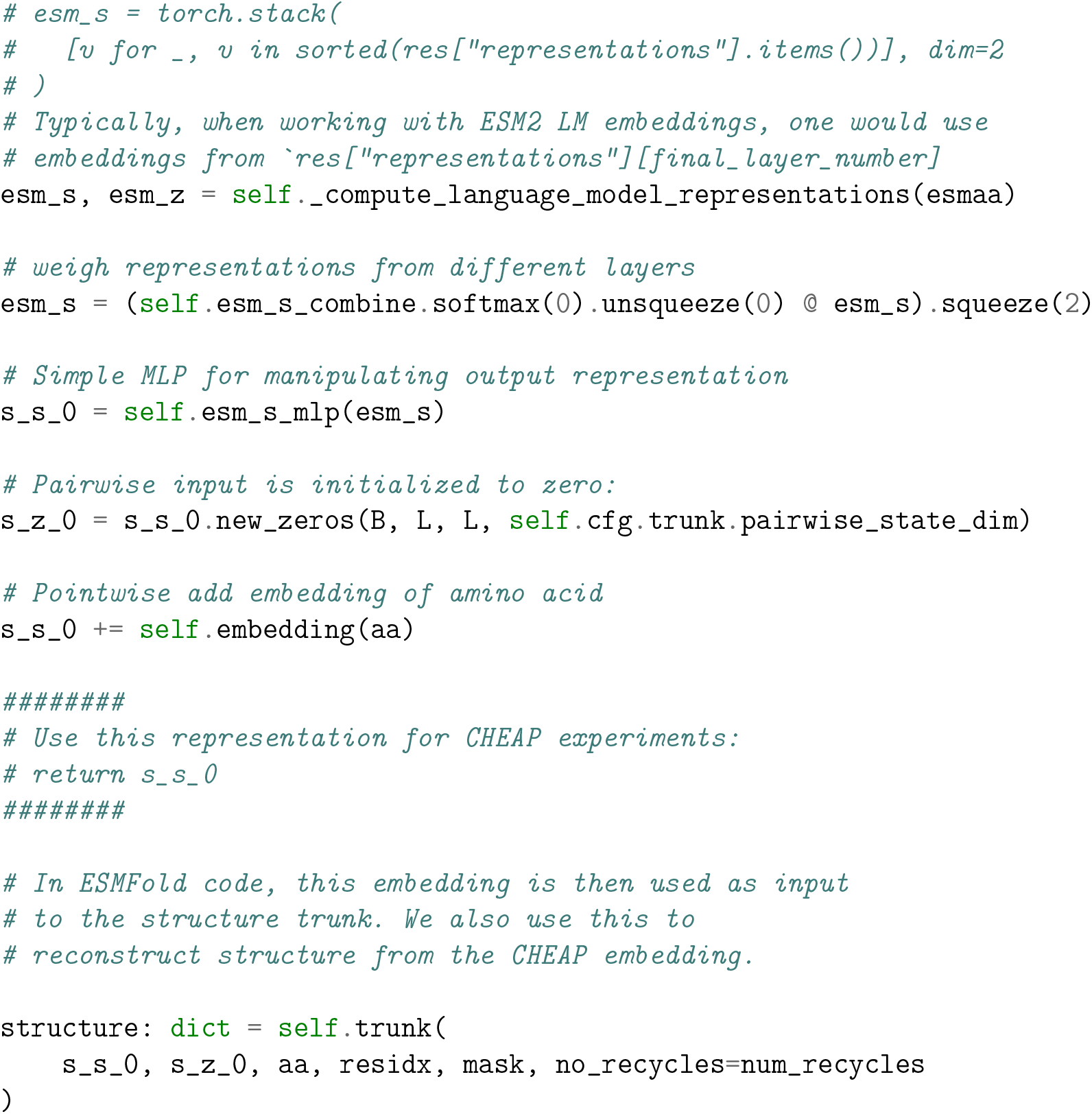

## Appendix B. FSQ codebook

For consistency when comparing with VQ-VAE and the original FSQ paper [26], we use the same FSQ levels:

| Target Size $\vert C\vert $ | $2^8$ | $2^{10}$ | $2^{12}$ | $2^{14}$ | $2^{16}$ |
| --- | --- | --- | --- | --- | --- |
| FSQ levels | [8, 6, 5] | [8, 5, 5, 5] | [7, 5, 5, 5, 5] | [8, 8, 8, 6, 5] | [8, 8, 8, 5, 5, 5] |

## Appendix C. Compression Results on Sequence-Only Datasets

TMScore is calculated with respect to the ESMFold predicted structure, rather than the ground truth structure.

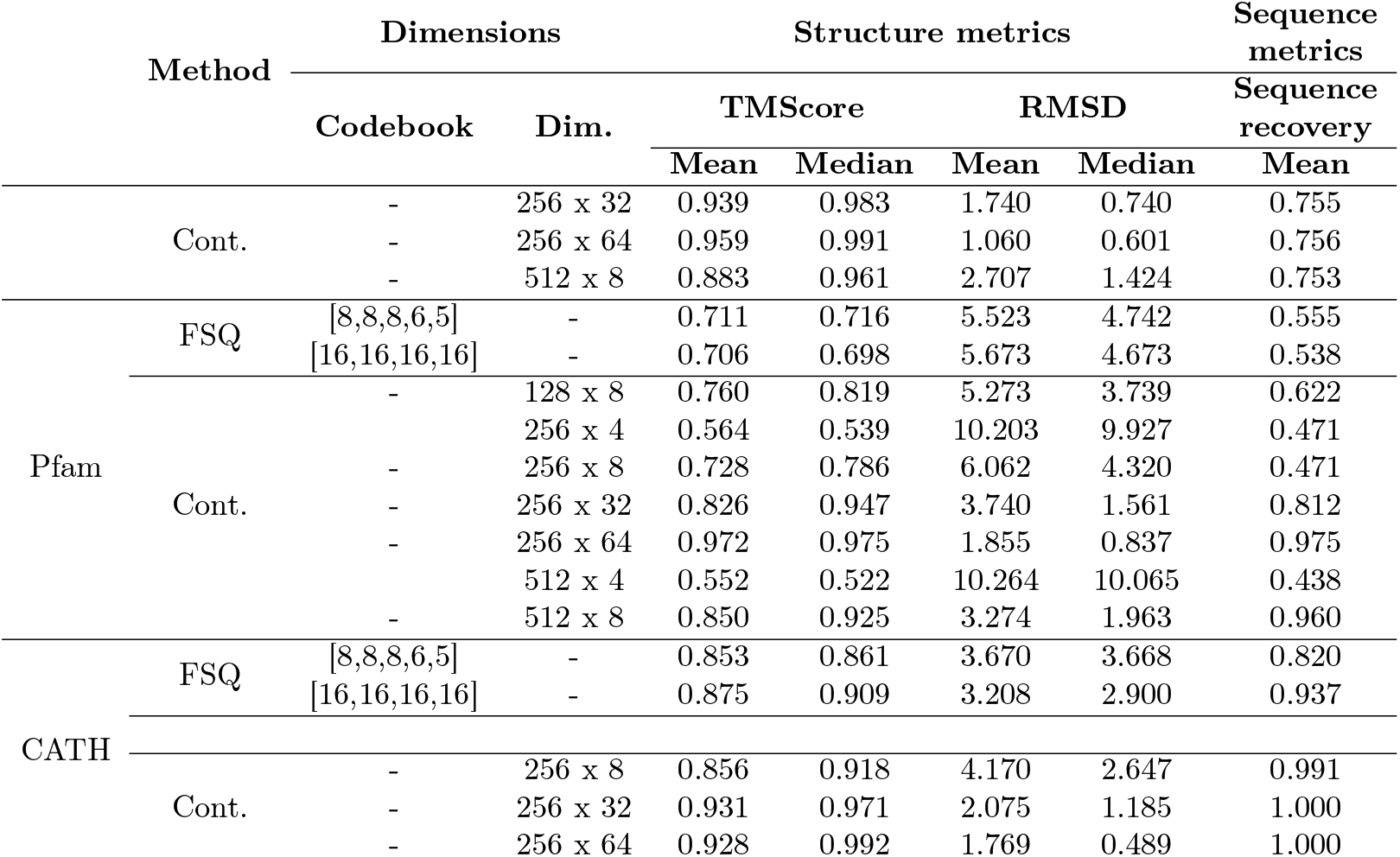

## Appendix D. Comparison with Other Embedding Methods on PEER Benchmark

Extending Figure 5, we also include comparisons to other reported performances in Xu et al. [45]:

**Table D.1:** Benchmarks on function and localization.

| | # Dimensions | Flu ↑ | Sta ↑ | β-lac ↑ | Sol ↑ | Sub ↑ | Bin ↑ |
| --- | --- | --- | --- | --- | --- | --- | --- |
| DDE | 400 | 0.64 | 0.65 | **0.62** | 0.60 | 0.49 | 0.77 |
| Moran | 240 | 0.40 | 0.32 | 0.38 | 0.58 | 0.31 | 0.56 |
| LSTM | 640 | 0.49 | 0.53 | 0.14 | 0.70 | 0.63 | 0.88 |
| Transformer | 512 | 0.64 | 0.65 | 0.26 | 0.70 | 0.56 | 0.76 |
| CNN | 21 | **0.68** | 0.64 | 0.78 | 0.64 | 0.59 | 0.83 |
| ResNet | 512 | 0.64 | 0.13 | 0.15 | 0.67 | 0.52 | 0.79 |
| ProtBert | 1024 | 0.34 | 0.70 | **0.62** | 0.59 | 0.59 | 0.82 |
| ESM-1b | 1280 | 0.43 | **0.75** | 0.53 | 0.67 | **0.80** | 0.92 |
| **CHEAP (ours)** | 4 | 0.14 | 0.40 | 0.13 | 0.60 | 0.33 | 0.68 |
|  | 8 | 0.22 | 0.44 | 0.17 | 0.64 | 0.45 | 0.74 |
|  | 16 | 0.27 | 0.55 | 0.23 | 0.65 | 0.54 | 0.84 |
|  | 32 | 0.28 | 0.56 | 0.28 | 0.67 | 0.57 | 0.87 |
|  | 64 | 0.31 | 0.56 | 0.28 | 0.69 | 0.62 | 0.90 |
|  | 128 | 0.41 | 0.58 | 0.38 | 0.70 | 0.68 | 0.90 |
|  | 256 | 0.47 | 0.60 | 0.41 | 0.71 | 0.72 | 0.92 |
|  | 512 | 0.51 | 0.63 | 0.36 | **0.72** | 0.74 | 0.93 |
| No compression | 1024 | 0.52 | 0.64 | 0.45 | **0.72** | 0.76 | **0.94** |

**Table D.2:** Benchmarks on structural and contact tasks.

|  | # Dimensions | Cont | Fold | SSP | Yst |
| --- | --- | --- | --- | --- | --- |
| DDE | 400 | — | 0.10 | — | 0.56 |
| Moran | 240 | — | 0.07 | — | 0.53 |
| LSTM | 640 | 0.26 | 0.08 | 0.69 | 0.54 |
| Transformer | 512 | 0.18 | 0.09 | 0.60 | 0.54 |
| CNN | 21 | 0.10 | 0.11 | 0.66 | 0.55 |
| ResNet | 512 | 0.20 | 0.09 | 0.70 | 0.49 |
| ProtBert | 1024 | 0.40 | 0.11 | 0.82 | 0.54 |
| ESM-1b | 1280 | 0.46 | 0.30 | 0.83 | **0.66** |
| **CHEAP (ours)** | 8 | 0.28 | 0.15 | 0.82 | 0.45 |
|  | 64 | **0.42** | 0.45 | 0.85 | 0.48 |
|  | 128 | 0.38 | 0.47 | 0.85 | 0.51 |
|  | 256 | 0.23 | 0.50 | 0.85 | 0.51 |
|  | 512 | 0.37 | **0.53** | **0.86** | 0.46 |

## Appendix E. Compressed Representation TSNE Plots

TSNE of embeddings after and before compression, colored by the “Architecture” level of the CATH classification hierarchy. Qualitative examination shows that despite 128× compression, fold and topology information is still large preserved.

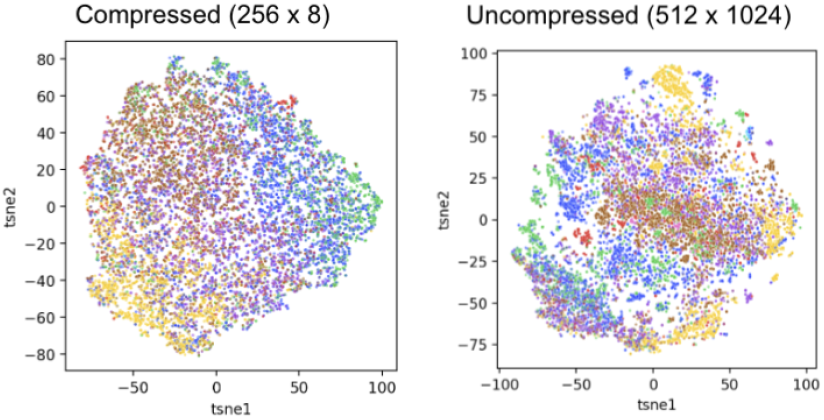

## Appendix F. Token Masking on Structure Reconstruction

Mirroring the analysis in Figure 7, we also examine the effect of “disturbing” the tokens on the output structure (Figure 4C), as a sanity check of how tokens encode structural information. The examined model uses FSQ levels [16, 16, 16, 16], and has an implied codebook size of *C* = 16^4^ = 65536. Since there is no `[MASK]` token in the vocabulary, we set different pre-quantization embeddings to 0. Generally, changing a subset of the tokens changes the structure, though since the codebook representations are not evenly spread through out, setting the first channels to zero has less of a detrimental effect.

**Figure F.8:**
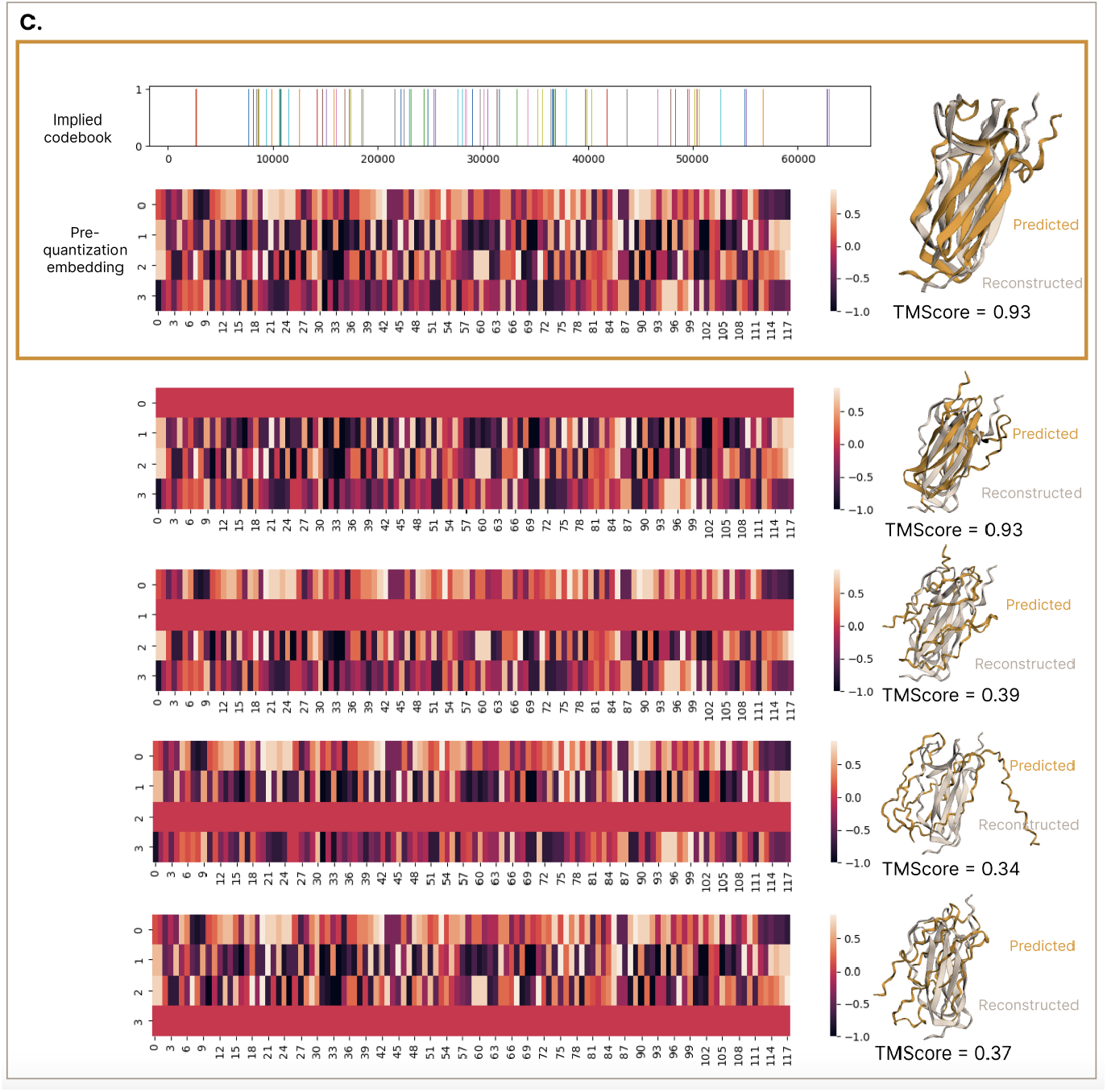
Exploring effects of token manipulation on the structure space for the [16, 16, 16, 16] FSQ model, by setting some values to 0 just prior to quantization. The yellow box (top) denotes the original pre-quantization embedding and discrete codes representations of all-atom structure.

## Footnotes

1 Note that the embedding dimension has already been reduced from the *L ×* 2560 ESM-3B embeddings.

